# Convergent genetic adaptation in human tumors developed under systemic hypoxia and in populations living at high altitudes

**DOI:** 10.1101/2024.06.10.594693

**Authors:** Carlota Arenillas, José Ruiz-Cantador, Lucía Celada, Bruna Calsina, Eduardo García-Galea, Debayan Datta, Roberta Fasani, Ana Belén Moreno-Cárdenas, Juan José Alba-Linares, Berta Miranda, Ángel M. Martínez-Montes, Cristina Álvarez-Escolá, Beatriz Lecumberri, Elvira Ana González García, Shahida K. Flores, Emmanuel Esquivel, Yanli Ding, Rita Maria Regojo Zapata, Jose Juan Pozo-Kreilinger, Carmela Iglesias, Trisha Dwight, Christopher Muir, Amelia Oleaga Alday, Maria Elvira Garrido-Lestache Rodríguez-Monte, Maria Jesús Del Cerro, Isaac Martínez-Bendayán, Delmar M. Lourenço, Maria Adelaide A. Pereira, Nelly Burnichon, Alexandre Buffet, Craig Broberg, Paxton Dickson, Mario Fernandez Fraga, José Luis Llorente Pendás, Joaquín Rueda Soriano, Francisco Buendía Fuentes, Sergio P.A. Toledo, Roderick Clifton-Bligh, Rodrigo Dienstmann, Jaume Capdevila, Anne-Paule Gimenez-Roqueplo, Judith Favier, Donate Weghorn, Paolo Nuciforo, William Young, Alexander R. Opotowsky, Anand Vaidya, Irina Bancos, Mercedes Robledo, Anna Casteràs, Laura Dos-Subirà, María Dolores Chiara, Igor Adameyko, Patricia L.M. Dahia, Rodrigo A. Toledo

## Abstract

*EPAS1*^HIF2α^ is the primary gene implicated in systemic hypoxia adaptation. Conversely, aberrantly activated *EPAS1*^HIF2α^ acts as a tumor driver against which anti-tumor therapeutics are proven effective. We elucidated connections between adaptation to systemic hypoxia in high-altitude populations, such as Tibetans and Sherpas, and human tumors. Similar to the accelerated adaptability observed in high-altitude populations via genetic introgression, tumors from patients with hypoxia since birth exhibited impaired DNA repair and increased mutation burden. As in high-altitude dwellers, *EPAS1*^HIF2α^ genetic variants were positively selected within sympathetic tumors developed under hypoxia, with a consistently high frequency of 90%. Bulk and single-cell RNA sequencing followed by *in vitro* studies have shown that hypoxia and *EPAS1*^HIF2α^ gain-of-function tumor mutations induce *COX4i2* expression and impair mitochondrial respiration, indicating that decreased cellular oxygen consumption may confer a proliferative advantage in hypoxia. Analyzing medical data from a patient cohort with hypoxia since birth who developed/did not develop tumors revealed tissue-specific and time-dependent tumorigenic effects of systemic hypoxia, which is limited to oxygen-sensitive and responsive cells, particularly during the postnatal period. This study supports connections between the *EPAS1*^HIF2α^ genetic adaptation in human tumors developed under systemic hypoxia to populations living in high altitudes. The genetic adaptations in populations to different stressors can be explored further to understand tumorigenesis and tumor evolution.

## Introduction

Atmospheric oxygen emerged approximately 2.3 billion years ago and triggered essential cellular, developmental, biochemical, and physiological adaptations, pivotal for survival and evolution of life^1^, of which ancestral bacteria (endosymbiotic *Alphaproteobacterium*^2^) integration allowed mitochondrial respiration of molecular oxygen (O_2_) through the electron transport chain (ETC), causing a 15-fold increase in ATP production compared to anaerobic metabolism. Mitochondria also play key roles in biomolecular anabolism and signal transduction via various mechanisms, including reactive oxygen species (ROS) production^3,4^. Increased energy availability and enhanced cellular communication have facilitated the emergence of multicellular animals, which require additional adaptations to effectively regulate and respond to fluctuating oxygen levels across different body regions^5^. In this context, the transcription factor hypoxia-inducible factor 1 (HIF1) is a key oxygen homeostasis regulator and primarily coordinates adaptive responses to fluctuating cellular oxygen levels^6^. During hypoxic events, specific prolines within the oxygen-dependent domain of HIF1α unit stop undergoing hydroxylation and degradation by VHL, leading to its stabilization and translocation to the nucleus, where it binds with the stable HIF1β unit^7–10^. This protein complex initiates the expression of numerous genes, including those associated with metabolic reprogramming (e.g., the Warburg effect) and neoangiogenesis, among other processes aimed at restoring oxygen homeostasis of the affected cells.

Throughout evolution, HIF2α, encoded by the *EPAS1* gene, emerged from the *HIF1α* gene duplication. HIF2α plays a versatile role in regulating and facilitating adaptations to whole body (systemic) hypoxia^11–13^. Specifically, HIF2α is required for embryonic development and organ function involved in O_2_ sensing and response to systemic hypoxia, such as the parasympathetic carotid body, sympathetic paraganglia, and adrenal medulla^14^. When mature, about a week after birth, these organs function as follows: first, glomus cells within the carotid body sense low O_2_ tension in the blood (hypoxemia); second, through afferent splanchnic nerves, the carotid body signals the sympathetic chromaffin cells of the adrenal medulla, which abundantly secrete adrenaline and noradrenaline (collectively known as catecholamines); third, the nervous stimulus combined with high adrenaline levels elicit a rapid compensatory response to hypoxia, involving hyperventilation, tachycardia, and increase venous tonus^15,16^. Nevertheless, the immature adrenal medulla from birth until day 7 of line, when it is innervated by the splanchnic nerves, can autonomously detect low oxygen levels, and secrete extremely high levels of catecholamines, such as norepinephrine, into the bloodstream in response to hypoxia, which is a crucial for neonatal adaptation to extrauterine life^17,18^.

Moreover, HIF2α is directly involved in adaptation to prolonged hypoxia. For example, when mountaineers climb high-altitude peaks and experience hypobaric hypoxia, HIF2α is activated in the kidney and augments erythropoietin (EPO) production, a hormone which stimulates erythropoiesis to enhance oxygen transport and tissue oxygenation^19–21^.

Consistent with *EPAS1*^HIF2α^ evolutionary specialization in systemic hypoxia adaptation, several studies have observed extreme positive selection of genetic *EPAS1* genetic variants in highlanders compared to lowlanders across many species^22–30^. *EPAS1* is indicated as a major target of genetic adaptation to high-altitude hypoxia. These highly advantageous *EPAS1*^HIF2α^ haplotypes in humans and other mammals present a loss-of-function effect in the HIF2α-EPO axis, avoiding excessive production of red-blood in hypoxia and augmenting blood perfusion to the organs and periphery ^31–34^. Beneficial *EPAS1*^HIF2α^ variants frequently originate from genetic introgression because of reproduction with a related species that is already adapted to hypoxic environments rather than occurring *de novo*^35,36^. For humans, the *EPAS1*^HIF2α^ haplotype likely originated via archaic introgression from the Denisovans who lived near the Tibetan plateau^36–38^, and is currently present in 90% of the highlander Tibetans and Sherpas. This Denisovan-like *EPAS1* haplotype is not found elsewhere, representing the strongest selection of any gene reported in humans^25,36,38^.

*EPAS1*^HIF2α^ has been identified as a *bona fide* oncogene, promoting tumorigenesis and tumor development^39–41^. The tumorigenic role of HIF2α was recently confirmed by the successful use of an on-target HIF2α inhibitor in hypoxia-driven tumors^42–46^. The oncogenic function of *EPAS1*^HIF2α^ is tissue-dependent and impacts cell lineages, wherein it controls embryonic development, such as the carotid body (derived from parasympathetic tissue), and sympathetic paraganglia and adrenal medulla^14^. Tumors that arise from these cells sensitive to *EPAS1*^HIF2α^ are classified as pheochromocytomas or paragangliomas, collectively referred to as PPGLs^40,47,48^. They are challenging for oncological management and characterized by rarity, therapeutic orphan status, absence of metastatic biomarkers, and a lack of suitable animal models. PPGLs are highly heritable neoplasms frequently driven by single, mutually exclusive germline or somatic mutations, which often directly or indirectly activate HIF2α^49,50^. Earlier studies suggested that environmental systemic hypoxia resulting from high-altitude environments in the Andes and Colorado^51–53^ or pathological cyanotic congenital heart disease (CCHD), could pose a risk for PPGL development ^54^. Epidemiological studies have estimated that patients with CCHD have a six-fold higher chance of developing PPGL than the general population or individuals with non-CCHD^54,55^.

The combination of systemic hypoxia, usually since birth, and increased risk of developing PPGL makes patients with CCHD unique human models for studying the role of hypoxia in tumorigenesis. Recent studies including those from our group have revealed that PPGLs from patients with systemic hypoxia, such as CCHD or sickle cell disease, harbor somatic gain-of-function mutations in the *EPAS1*^HIF2α^ gene^56–59;^ however, their mechanistic insights remain largely unknown.

In this study, we explored genetic, *in vitro*, population, and clinical data to determine the mechanism time of occurrence, and reason for *EPAS1*^HIF2α^ mutation generation in tumors developed under systemic hypoxia. We identified significant molecular convergence between human tumors from patients with chronic systemic hypoxia and those adapted to high-altitude hypoxia, such as Tibetans and Sherpas. Our study reveals a previously unrecognized level of convergence where natural populations and tumor cells subjected to similar environmental stressors, such as hypoxia, develop analogous genetic and molecular adaptations. This discovery could guide future studies on the links between natural adaptation and tumorigenesis, paving the way for the identification of new tumor drivers and therapeutic vulnerabilities.

## Results

### Highly frequent EPAS1^HIF2α^ clonal mutations in sympathetic PPGL tumors developed under systemic hypoxia

We assembled and examined a large cohort of 34 PPGL tumor samples collected from 27 patients with CCHD across five distinct countries (**Fig. S1, S2, table S1**). We investigated the *EPAS1*^HIF2α^ status, potential genotype-phenotype correlations, and underlying molecular mechanisms of tumor adaptation and survival under systemic hypoxia. Most tumors (27/34, 79.4%) were sympathetic catecholamine-secreting PPGLs, comprising pheochromocytomas and abdominal and thoracic paragangliomas. Targeted Sanger sequencing revealed *EPAS1*^HIF2a^ mutations in 24/27 sympathetic CCHD-PPGLs (88.9%). All mutations occurred exclusively in the tumor DNA and not in the germline DNA, and clustered within the oxygen-dependent degradation domain of HIF2α^60^ and were missense alterations (L529P, A530P/T/V, P531A/R/L/S, Y532C, L542R/P) except one in-frame deletion (P534_N536del) **(table S1)**. A total of 12/14 (85.7%) abdominal and thoracic paragangliomas and 12/13 (92.3%) pheochromocytomas were mutated (**Fig. 1A**). Two patients had multiple tumors and all of them carried a distinct *EPAS1* missense somatic mutation (P16 and P21, **Fig. 1B**). Whole-exome sequencing (WES) excluded germline or somatic mutations in other known PPGL susceptibility genes, emphasizing the significance of *EPAS1* mutations as CCHD-PPGL tumor genetic driver. These results highlight a strong convergent evolution of *EPAS1* in PPGL tumorigenesis under systemic hypoxia at the inter and intra-patient levels.

**Fig. 1.**
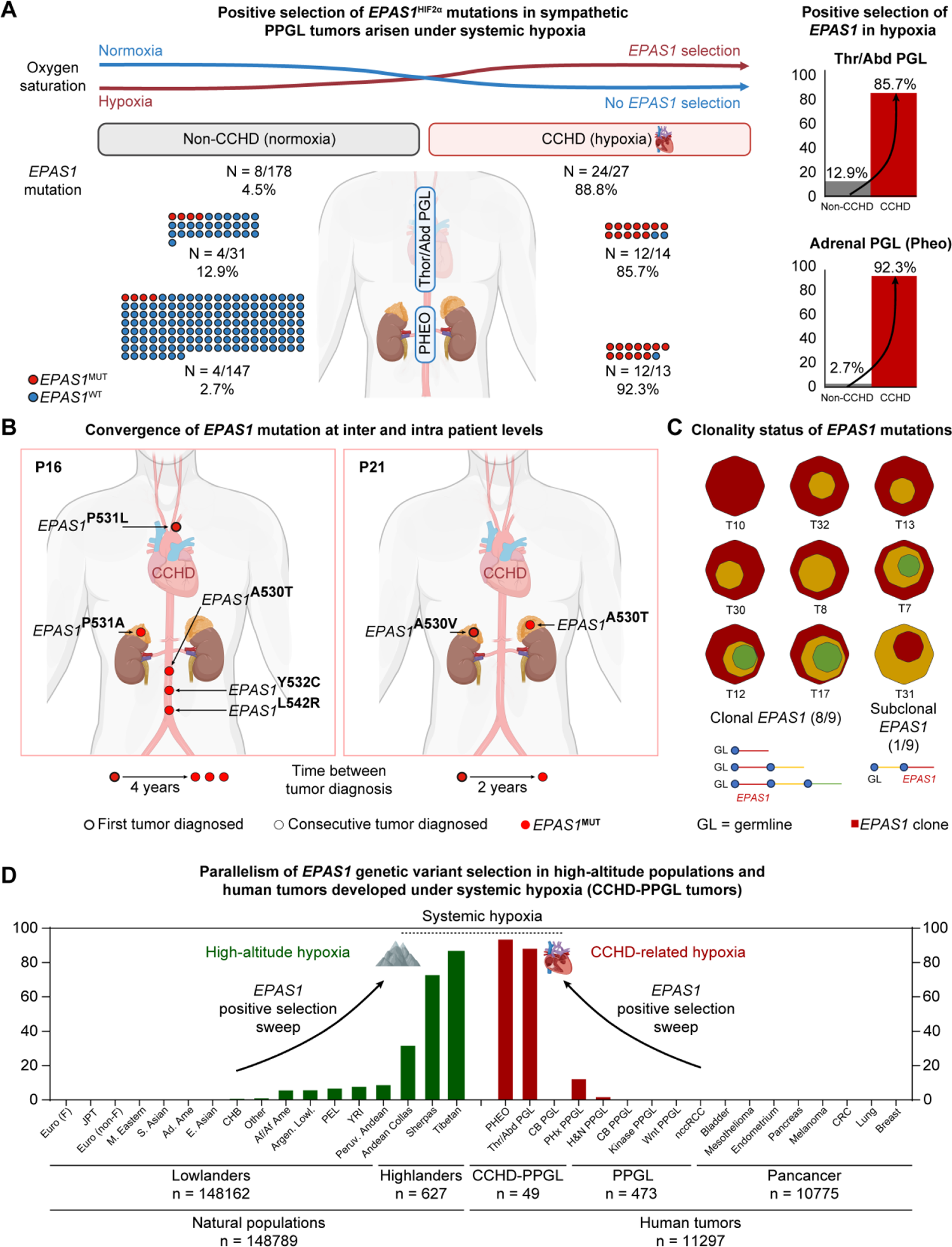
**A)** Positive selection of *EPAS1* mutations in our cohort of PPGL tumors from patients born with CCHD and systemic hypoxia (24/27, 88.8%, right side) compared to those without (non-CCHD cohort from the TCGA, 8/178, 4.5%, left side). Red circle: *EPAS1*^MUT^ (*EPAS1*-mutated); blue circle: *EPAS1*^WT^ (*EPAS1*-wild-type). Considering tumor location, *EPAS1* mutations were highly enriched in thoracic and abdominal paraganglioma (PGL) from CCHD patients compared to non-CCHD (85.7vs. 12.9%) and in pheochromocytomas (Pheo) from CCHD patients compared to non-CCHD (92.3% vs 2.7%). **B)** *EPAS1* mutational convergence in patients with multiple primary CCHD-PPGLs. Patient 16 (left) developed five independent primary tumors. The first PGL tumor was diagnosed in the mediastinum at 29 years of age; four years later the patient developed three peri-aortal paragangliomas and a right pheochromocytoma, each carrying a different *EPAS1* missense mutation (A530T, P531A, P531L, Y532C, L542R). Patient 21 (right) developed metachronous bilateral pheochromocytoma (the first at 32 years of age) within a two-year period and each tumor carried a different *EPAS1* mutation (named A530V and A530T). Patient 1 (not shown) presented with an *EPAS1*-wt carotid body and two months later was diagnosed with a left pheochromocytoma that harbored an *EPAS1* mutation (P531S). Red circle: *EPAS1*^MUT^; bold outline circle: first tumor diagnosed; black outline circle: consecutive tumors diagnosed. **C)** Clonality analysis of *EPAS1* mutations in CCHD-PPGL tumors: 88% (8/9) tumors harbored an *EPAS1* clonal mutation with a median cancer cell fraction (CCF) of 1, **tables S3-S5**. The clone where *EPAS1* mutation is included is represented in red. **D)** Allele frequencies of *EPAS1* genetic variants for different populations living in high and low altitude regions (left side, in green) and for different tumor type cohorts from patients with normoxia from publicly available cancer genomics databases and for PPGL tumors from patients with CCHD developed under hypoxia (right side, in red color). The figure displays a representation of the entire cohort of individuals and tumor samples analyzed, listed in table S6. Functional *EPAS1* genetic variants are extremely positively selected exclusively in natural populations genetically adapted to high-altitude hypoxia (up to 87% in Tibetans) and in sympathetic PPGL tumors from patients with CCHD born with hypoxia (up to 90%). In contrast, *EPAS1* variants are extremely rare and mostly absent in all low-lander populations and in cancer cohorts from patients with normoxia; non-CCHD sympathetic PPGLs harboring up to 4.5% of *EPAS1* mutations were the exception. These results strongly support systemic hypoxia as a major driver of *EPAS1* genetic variant positive selection in natural populations and in human tumors (see table S6). Admixed American (Ad. Ame); African/American (Af./Am.); Argentinian Lowlanders (Argen. Lowl.); Carotid Body (CB); Colorectal Cancer (CRC); Cyanotic congenital heart disease (CCHD); East Asian (E. Asian); Finnish (F); Head and Neck (H&N); Japanese in Tokyo (JPT); Non-clear-cell renal cell carcinomas (nccRCC); Peruvians residing in Lima (PEL); Pheochromocytomas and paragangliomas (PPGL); Pseudohypoxia (PHx); South American (S. American); South Asian (S. Asian); Thoracic/Abdominal (Thr/Abd); Yoruba from Ibadan (YRI).

Only 8/178 (4.5%) PPGLs from patients from the TCGA project, without CCHD^61^, carried an *EPAS1* mutation, indicating a 20-fold increase in *EPAS1* mutation frequency in patients with hypoxia vs. normoxia (89% vs. 4.5%, p < 0.0001) (**Fig. 1A**). Selection inference confirmed that *EPAS1* was the only gene under statistically significant and strong positive selection in our CCHD-PPGL cohort, with dN/dS=702 (q=0, **table S2**). When further stratifying by tissue type, the inferred selection strength increases to dN/dS=926 in the sympathetic sub-cohort, while dN/dS=0 in the parasympathetic sub-cohort, as all *EPAS1* mutations were found in sympathetic PPGLs. In PPGL tumors from non-CCHD patients of the TCGA, *EPAS1* was the gene with the third-largest signal of positive selection (after *HRAS* and *NF1*) and marginally significant (dN/dS=110, q=0.1, **table S2**). This contrasts with the rest of the TCGA cohort, in which *EPAS1* is not selected, neither at the pan-cancer level nor in any of the individual cancer types.

Furthermore, we assessed the neoplastic cell proportion of *EPAS1*-mutated (*EPAS1*^MUT^) clones in PPGL tumors developed in normoxia (TCGA cohort) and in systemic hypoxia (CCHD cohort) via calculating the cancer cell fraction (CCF)^62^. *EPAS1* mutations presented extremely high CCF levels in both cohorts (median of 1 and 0.98), a result that clearly supports *EPAS1* mutations as initial genetic events in PPGLs (**Fig. 1C**, **fig. S3, S4, table S3-S5**).

### EPAS1^HIF2a^ mutation absence in parasympathetic PPGL tumors and across tumor types

Contrasting the extreme high and moderate *EPAS1* mutation frequency in sympathetic PPGL tumors from patients with CCHD (24/27, 88.8%) and in sympathetic PPGL tumors from patients without CCHD (TCGA) (8/178, 4.5%), respectively, targeted sequencing identified no *EPAS1* mutations in seven parasympathetic carotid body PPGLs from patients with CCHD (0/7) (**fig. S5**). WES of four of these tumors showed no mutations in complete coding regions of *EPAS1* gene, its homologue genes *HIF1α* and *HIF3α,* and any known PPGL-related pseudohypoxia genes. No *EPAS1* mutations were detected in additional parasympathetic PPGLs from patients without CCHD: 0/52 from a previous cohort^63^, and 0/214 from an unpublished cohort we collected (see methods^64^, p < 0.00001, **fig. S5**). Furthermore, *EPAS1* mutations were extremely rare across 33 human cancer types from the 212 studies available on cBioPortal^65^ (5/69,045 tumor samples, 0.007%) and TCGA Pancancer Atlas Studies^66^ (0/10,775 tumor samples, 0%) (**table S1 and S6)**. These results suggest that *EPAS1* somatic mutations occur exclusively in PPGL tumors of sympathoadrenal lineage and are 20-fold enriched in CCHD (systemic hypoxia) (**Fig. 1D**).

### EPAS1 evolutionary trajectory in systemic hypoxia-developed PPGL tumors parallels with that observed in highlander Tibetans and Sherpas

*EPAS1* genetic variants are rarely selected from numerous samples sequenced from global populations living at low altitudes, or from tumor samples across more than 30 tumor types. In contrast, we detected *EPAS1* mutations in 89% sympathetic PPGL tumors in patients with CCHD, paralleling the 90% *EPAS1* variant prevalence reported in Tibetans and Sherpas in the Himalayas ^23,25,36,67^ (**Fig. 1D**). This level of extreme positive selection and genetic variant fixation is rare in natural populations. For Sherpas, *EPAS1* variants were likely introgressed by archaic Denisovans to early modern human populations^36^, probably when both species co-inhabited the Tibetan plateau 30–45 thousand years ago^68–70^. Introgression and selection sweep of advantageous *EPAS1* genetic variants has been reported in different species and animal populations living at high altitudes^26–30^ (**Fig. 1D, fig. S6, table S6**) and represents the most common and efficient hypoxia adaptation method. The positive selection of introgressed *EPAS1* genetic variants in Himalayan populations indicates that evolutionary introgression is an accelerated evolution mechanism that greatly facilitates hypoxia adaptation (**fig. S7**). Other human populations living at similar altitudes as the Tibetans, such as Andean Collas from Argentina and Peruvian Andean (living in altitudes up to 4,450m), present marked maladaptation features, that is, high-altitude or chronic mountain sickness^71,72^. The high Andes was inhabited more recently than the Tibet plateau (approximately 10 thousand vs. 40 thousand years ago) and without indicating genetic introgression^73–76^. Therefore, both the Andean Collas and Peruvian Andean populations harbor lower *EPAS1* genetic variants than the Tibetans (9% vs. 32% vs. 90%, respectively^73–76)^ (**Fig. 1D, table S6-S7**).

The evolutionary parallelism observed in high-altitude dwellers and in the PPGL tumors from patients with CCHD experiencing systemic hypoxia since birth is unique. In natural populations, this phenomenon is propelled by accelerated adaptation and characterized by introgression and selection sweeps^36,38,69,77^. Likewise, we identified and characterized that CCHD-PPGL tumors developed under systemic hypoxia likely harbored an accelerated process that enhanced hypoxia adaptation, as follows.

### Faulty DNA repair and increased mutation burden in systemic hypoxia-developed tumors

Previous studies showed decreased DNA repair, increased mutability, and increased adaptability in tumor cell lines cultivated under hypoxic stress^78–84^ and increased tumor mutation burden in tumors with increased intratumoral hypoxia, measured by RNAseq-based signatures^78,79^; hence, we hypothesized that similar processes could be driven by systemic hypoxia. Our results showed that non-neoplastic tissues adjacent to tumors stained positive for microsatellite instability (MSI) proteins, which was expected; however, most *EPAS1*^MUT^ sympathetic PPGL tumors from patients with hypoxic CCHD displayed faulty DNA repair system proteins (**fig. S8**). Specifically, 7/9 (77%) tumors available for immunohistochemistry staining presented evidence of MSI, characterized by the expression loss of mismatch repair proteins, such as MLH1, MSH2, MSH6, and PMS2 (**Fig. 2A**), which indicated reduced DNA repair function. On the contrary, all four *EPAS1*-wild-type (*EPAS1*^WT^) parasympathetic CCHD-PPGL tumors expressed MSI proteins MLH1, PMS2, MSH2, and MSH6 (**Fig. 2B**). Differences in MSI statuses between tumors from sympathetic and parasympathetic lineages are clear, 77% vs. 0%, and semi-quantitative H-score quantification shown in **Fig. 2C**. MSI generally occurs in 3–4% human tumors (exception being colorectal tumors with 20% MSI^85^). Therefore, the link between systemic hypoxia and MSI status we found in sympathetic CCHD-PPGLs is noteworthy. We generated WES data of nine sympathetic CCHD-PPGL tumors developed under hypoxia from our cohort and compared them with 118 sympathetic PPGL tumor samples from patients without CCHD from the TCGA cohort^61^. We observed a significantly higher average MSI genomic score, global copy number alterations, and tumor mutation in the sympathetic CCHD-PPGL tumors developed under hypoxia (average of 15 somatic mutations vs. 9 somatic mutations, p = 0.02, **Fig. 2D–F, table S8**). These results suggest a process of accelerated adaptability within sympathetic CCHD-PPGL tumors developed under hypoxia via hampering DNA repair and increasing the genetic instability and mutation pool within the tumor, which may favor the occurrence of *EPAS1* mutations that are then strongly selected during hypoxia (**fig. S9**).

**Fig. 2.**
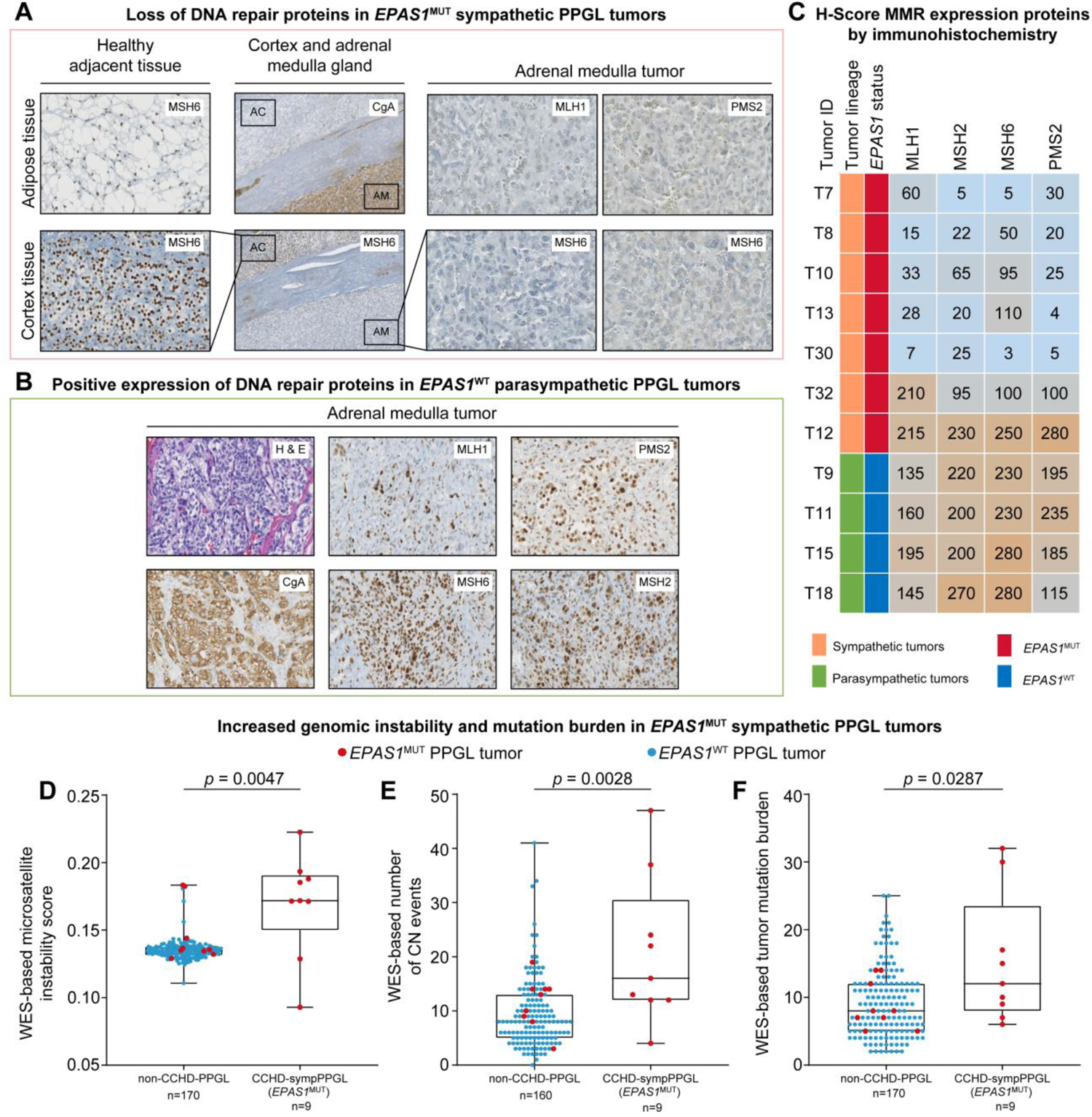
**A)** Immunohistochemistry staining showing decreased expression of DNA repair machinery proteins observed in most *EPAS1*^MUT^ sympathetic PPGL tumors from patients with CCHD (7/9, 77,7%). Results of *EPAS1*^MUT^ sympathetic PPGL tumor T7 are shown: i) expected positive expression of the MSH6 microsatellite instability (MSI) biomarker in PPGL tumor-surrounding tissues from patients with CCHD, such as adipose and adrenal cortical (40× magnification); ii) positive expression of chromogranin A in the adrenal medulla (PPGL tumor) and not cortex; iii) positive expression of MSH6 in the adrenal cortex but not in the PPGL tumor (10× magnification), and iv) decreased expression of MSI biomarkers such as MLH1, MSH2, MSH6, and PMS2 in the CCHD-PPGL neoplastic cells (40× magnification). **B)** When compared with PPGL parasympathetic tumor (T18) with *EPAS1*^WT^ status, all MSI biomarkers show a positive expression in the non-neoplastic and neoplastic cells. Hematoxylin and eosin and chromogranin A are also indicated as positive controls. **C)** Immunohistochemistry from MSI biomarkers in all available tumor samples, scored semi-quantitatively using an H-score for nucleus. Sympathetic tumors have a diminished H-Score (below 100) (6/7, 86%), whereas all parasympathetic tumors present higher H-score, indicating differences of mismatch repair protein (MMR) expression between tumor linages. **D, E, F)** Germline:tumor paired whole-exome sequencing (WES) data was used to compare MSI Score, Copy Number Alterations (CNA) Burden (number of CN events), and Tumor Mutation Burden (TMB) between sympathetic PPGL tumors from patients without and with CCHD (N=170, left, and N=9, right, respectively)^61,122^. Red circle: *EPAS1*^MUT^ tumors and blue circle: *EPAS1*^WT^. Cyanotic congenital heart disease (CCHD); Pheochromocytomas and paragangliomas (PPGL).

### Mitochondrial respiration modulation as a potential mechanism for hypoxia adaptation driven by EPAS1 mutations in systemic hypoxia tumors

We investigated the molecular foundation underlying the extreme *EPAS1* mutation selection in sympathetic CCHD-PPGL tumors developed under hypoxia. Previous studies from our group and others showed that *EPAS1* missense genetic variants in PPGLs were gain-of-function mutations and promoted tumor formation *in vivo* via hampering Von Hippel-Lindau (VHL)-dependent proteasomal HIF2α degradation^40,48^. To identify *EPAS1-*HIF2α-target gene(s) involved in systemic hypoxia adaptation amongst numerous genes transcriptionally controlled by HIF2α, we compared data from bulk RNAseq-based transcription profiles of *EPAS1*^WT^ (n = 46) and *EPAS1*^MUT^ (n = 8) PPGL tumors from the TCGA cohort^61^ and *EPAS1*^WT^ (n = 11) and *EPAS1*^MUT^ PPGL tumors (n = 19) from a cohort with hemoglobin disorders and putative chronic systemic hypoxia^59^. Also, new transcription data profiles of the PC12 rat PPGL-derived cell line cultured at different timepoints (12, 24, 48 h and 36 days) under normoxic (n = 12) and hypoxic (n = 12) conditions (1% O_2_) were included in the analyses (**Fig. 3A, table S9-S11**).

**Fig. 3.**
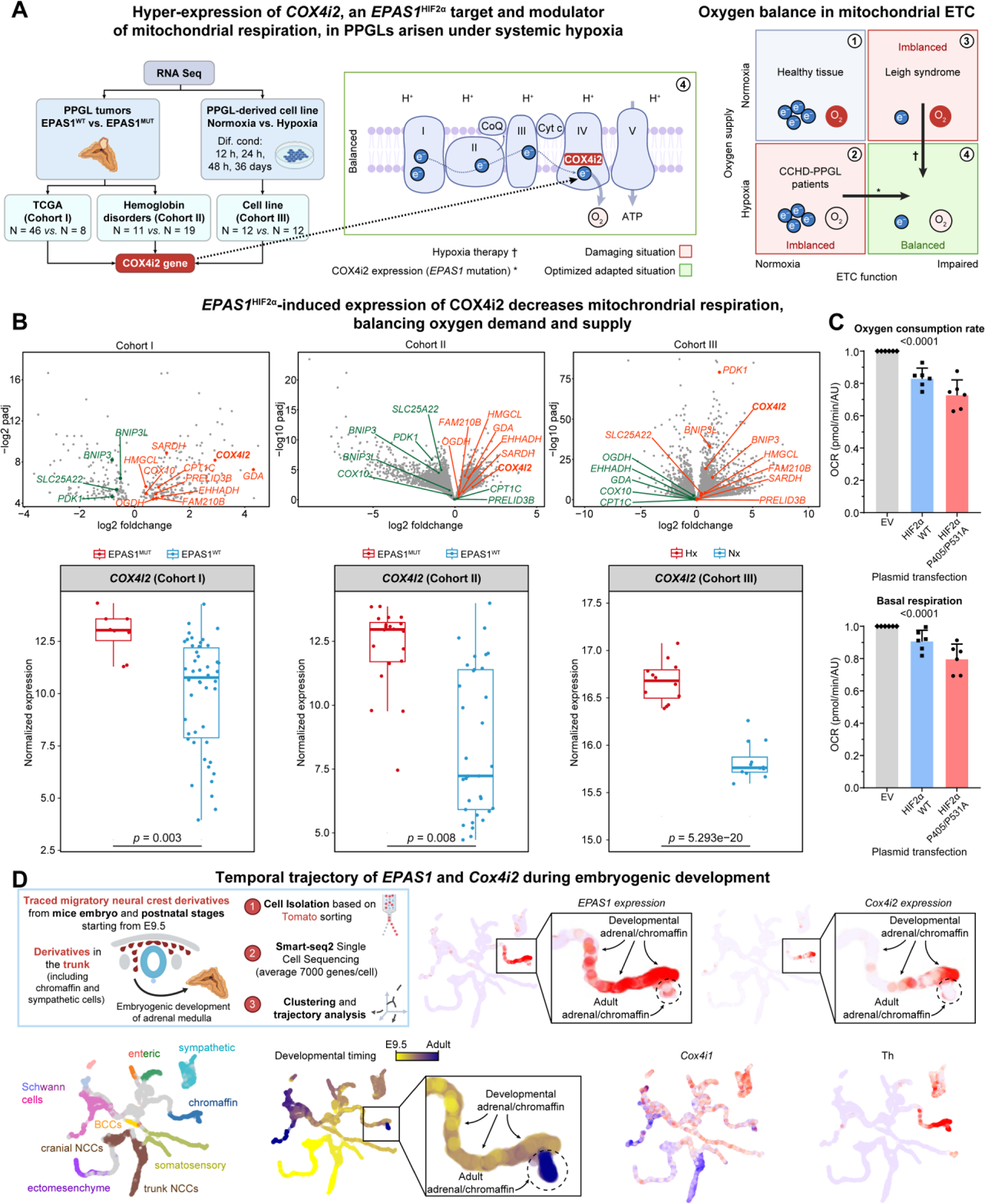
Transcription analyses for functional and developmental insights of *EPAS1* in PPGL tumorigenesis. **A)** Workflow of bulk RNA-Seq data analysis using three different cohorts: cohort I comprised non-CCHD sympathetic PPGL tumors wild-type (n = 46) or mutated (n = 8) for *EPAS1*^61,65^; cohort II comprised sympathetic PPGL tumors from patients with hemoglobin disorders: 11 tumors *EPAS1*^WT^ and 19 tumors *EPAS1*^MUT59^; cohort III comprised PPGL-derived cell line we cultured under normoxic and hypoxic conditions for 12h, 24h, 48h, and 36 days (12 replicates for normoxia and 12 for hypoxia). Of the 354 genes differentially expressed in patients with PPGL tumors, 14 mitochondrial genes were found, of which one was *COX4i2*, the less common isoform of COX (mitochondrial respiratory complex 4) and the terminal electron acceptor of the oxidative phosphorylation system. **B)** Upper part: Volcano plots of the differentially expressed genes between *EPAS1*^MUT^ and *EPAS1*^WT^ tumor samples from Cohorts I and II, and between hypoxia and normoxia conditions in Cohort III. Fourteen differentially expressed mitochondrial genes are highlighted (upregulated genes in orange color, downregulated genes in green color). Bottom part: Normalized expression of *COX4i2* in Cohorts I, II, and III. **C)** Mitochondrial respiration using Seahorse equipment (n = 6/group; mean ± SEM) in stable cell line HEK293T expressing Empty Vector (EV)-Control, HIF2α^WT^, and mutant HIF2α^P405A/P531A^ through Plasmid Transfection. Oxygen Consumption Respiration (OCR) (top), Basal Respiration (bottom) **D**) Single-cell RNA sequencing (scRNA-seq) data comprising the entire tree of neural crest lineage development, which includes differentiation trajectory from the neural crest towards chromaffin cells via Schwann cell precursor and Bridge state intermediates^89^. The scRNA-seq data shows that *EPAS1* and *COX4i2* expression is upregulated during development and downregulated in postnatal chromaffin/adrenal medulla cells, after the onset of breathing. Cyanotic congenital heart disease (CCHD); Pheochromocytomas and paragangliomas (PPGL).

From these three cohorts, we found significant hyperexpression of two genes that encoded mitochondrial proteins: *HMGCL*, involved in leucine and fatty acid metabolism^86^, and *COX4i2* (**Fig. 3A, 3B**). *COX4i2* warranted further evaluation as cytochrome c oxidase 4 (COX4) enzymes are final components of mitochondrial electron transport chain (ETC) that directly transferred electrons to oxygen and produced important cellular signaling intermediates, such as NADH, ROS, water, and ATP^87^. Fukuda *et al.* showed that cells cultivated under hypoxia switched expression of ubiquitous *COX4i1* to atypical *COX4i2* unit, which has a lower oxygen affinity, thus altering mitochondrial respiration under low oxygen conditions^88^ (**fig. S10**). We evaluated *EPAS1* mutation modulation of cellular aerobic respiration. Using the Seahorse respiratory assay, we found that HEK293 cell line that stably expressed *EPAS1* gain-of-function mutation (HIF2α P405/P531A) and induced *COX4i2* expression displayed reduced oxygen consumption rate compared to their *EPAS1*^WT^ expressing (HIF2α WT) counterparts [average 0.73 pmol/min/AU (range 0.63–0.87) vs average 0.83 pmol/min/AU (range 0.74-0.92), respectively (p < 0.0001)] and at basal respiration, an average 0.80 pmol/min/AU (range 0.69–0.91) in HIF2α P405/P531A and 0.91 pmol/min/AU (range 0.99–0.81) in HIF2α WT (p < 0.0001), as shown in **Fig. 3C**. These results indicate mitochondrial respiration modulation and cellular oxygen consumption optimization via *COX4i2* regulation as the functional molecular mechanism for strong positive *EPAS1* mutation selection in PPGLs generated under low-oxygen conditions.

### Temporal trajectory of EPAS1 and COX4i2 during embryogenic development and in EPAS1-mutated PPGL tumors

PPGLs are neural crest-derived tumors^50^. To evaluate the link between *EPAS1*^HIF2α^ and *COX4i2* and its temporal expression dynamics, we analyzed single-cell RNA sequencing (scRNA-seq) data comprising the entire neural crest lineage development tree, which included a differentiation trajectory from the neural crest towards chromaffin cells via Schwann cell precursor and Bridge state intermediates^89^ (**Fig. 3D**). We found that *EPAS1* and *COX4i2* expressions were correlated in an immature pre-birth chromaffin cell population, suggesting a common regulatory link that coordinated *EPAS1* and *COX4i2* in similar cells within the same time window in oxygen-sensitive and responsive neural crest-derived cell types (**Fig. 3D**). These scRNA-seq data highlight key roles of *EPAS1* and *COX4i2* in developing sympathetic systems; they are consistent with the mid-gestational lethality of *EPAS1* knockout mice caused by reduced catecholamine levels causing impaired cardiac function and mitochondrial homeostasis^14,90^. The concordance between scRNA-seq and genetic manipulation underscores the importance of *EPAS1* and *COX4i2* in orchestrating essential processes in immature chromaffin cells for proper sympathetic system development.

Our scRNA-seq data from in postnatal chromaffin/adrenal medulla cells after the onset of breathing showed that *EPAS1* and *COX4i2* expressions were downregulated at this later timepoint (**Fig. 3D**). This suggests a crucial developmental time window for *EPAS1* and *COX4i2,* which is weakened in mature chromaffin/adrenal medulla cells. We observed that *EPAS1*^MUT^ PPGL tumors showed a significant resurgence in *EPAS1* and *COX4i2* gene expression, suggesting a fetal-like transcriptional pattern in tumor chromaffin cells (**Fig. 3B, 3D**). *EPAS1* and *COX4i2* hyperexpression was confirmed in independent *EPAS1*^MUT^ PPGL tumor cohorts^59,91^ and was not observed in *EPAS1*^WT^ PPGL tumors, suggesting that it was driven by *EPAS1*-mutation-dependent HIF2α activation (**Fig. 3B**).

Integrating genetic, scRNA-seq, and bulk RNA data helped delineate a temporal window during embryological development, wherein *EPAS1* and *COX4i2* exhibited coordinated expression in immature chromaffin cells. This coordination diminishes postnatally upon cell maturation but resurfaces during *EPAS1* mutation-driven tumorigenesis.

### Temporal and tissue-specific impact of systemic hypoxia in tumorigenesis

Through an International Consortium, we built and curated a large dataset comprising 2,588 patients, including 1,599 patients diagnosed with CCHD without PPGL; 840 patients with PPGL without CCHD (tumors developed under normoxia); 149 patients with CCHD plus PPGL (tumors developed under hypoxia) (**Fig. 4A**). PPGLs were the only tumors enriched in CCHD, consistent with the findings of previous epidemiological studies^54,55^. We collected and analyzed clinical data regarding a) heart disease, such as CCHD type and heart surgery; b) hypoxia exposure, such as the timing of hypoxia initiation and duration, and O_2_ saturation (SatO_2_); c) PPGLs, such as age at diagnosis, tumor type, tumor multiplicity, metastatic status, and biochemical and genotype profiles.

**Fig. 4.**
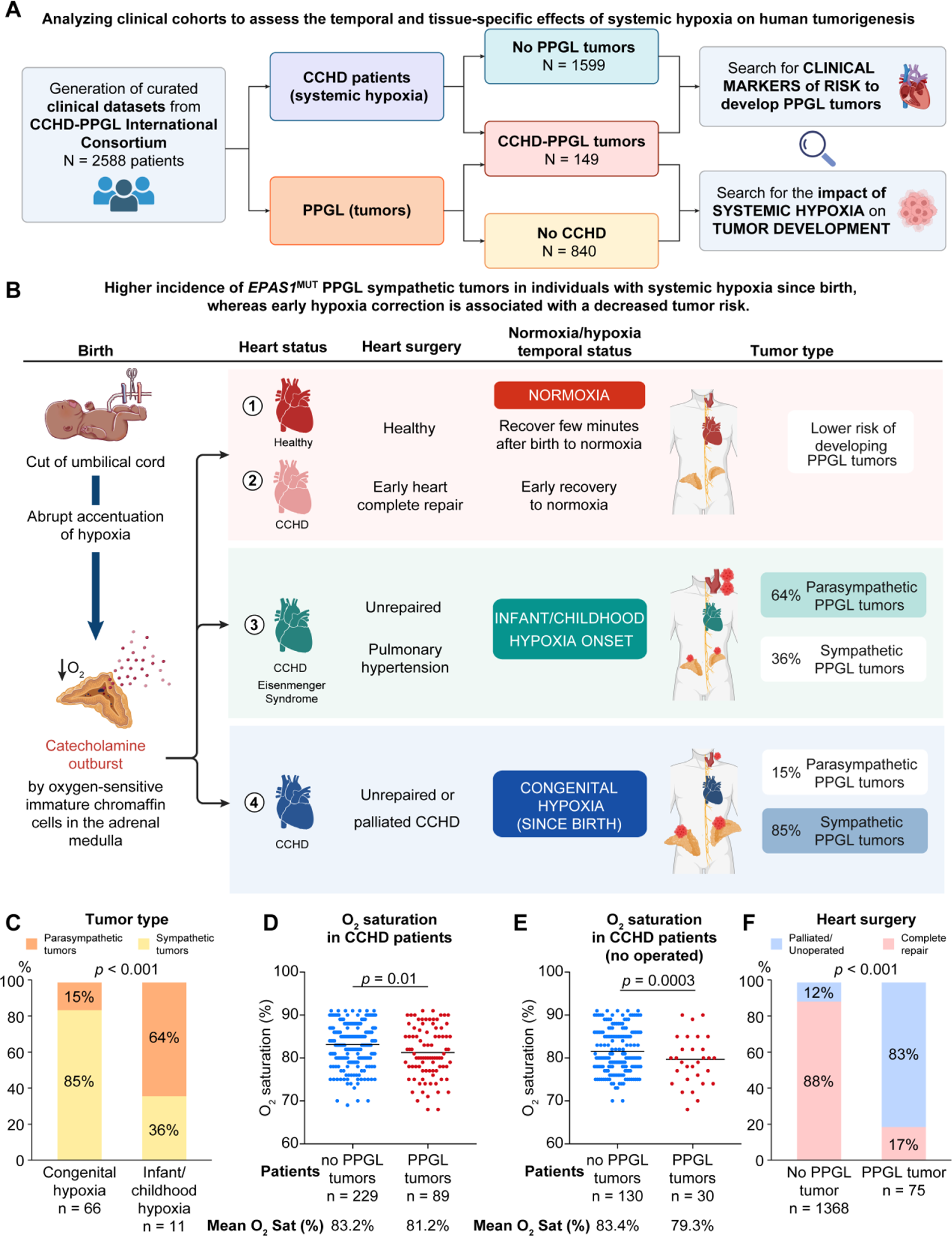
**A)** Overview of clinical datasets collected from the CCHD-PPGL International Consortium for the study. Clinical features from 2,588 patients have been collected and curated, involving patients with CCHD without PPGL tumors (N = 1,599), patients with CCHD-PPGL (N = 149), and patients with PPGL without CCHD (N = 840). Analyzing patients with CCHD with or without PPGL tumors (with systemic hypoxia in a determinate moment of their life), involved investigating clinical markers for the risk of developing PPGL tumors. Clinical characteristics from patients with PPGL tumors with or without CCHD were interrogated to determine the impact of systemic hypoxia on tumor development. **B)** Schematic figure depicting the hypoxia levels of patients with CCHD along their life and their catecholaminergic system. Depending on their systemic hypoxia, they are more prone to develop PPGL tumors. **C)** PPGL tumor types depending on the hypoxia exposure time. **D)** Oxygen saturation (%) in patients with CCHD with or without PPGL tumors. **E)** Oxygen saturation (%) in non-operated patients with CCHD with and without PPGL tumors. **F)** CCHD surgical management type and occurrence of PPGL tumors. Cyanotic congenital heart disease (CCHD); Pheochromocytomas and paragangliomas (PPGL).

To identify a potential time window for systemic hypoxia effects, we analyzed clinical outcomes of two patient groups with CCHD with PPGLs: one with severe heart complications causing hypoxia since birth and the other with Eisenmenger Syndrome, a rare heart defect that causes blood vessel damage, pulmonary hypertension, and systemic hypoxia that can start not right after birth, but during childhood or early adulthood^92^. In the first group, 85% (56/66) patients with hypoxia since birth developed catecholamine-secreting PPGL tumors originating in sympathetic system cells, such as the adrenal medulla and/or paraganglia (**Fig. 4B, 4C, table S12**). The remaining 15% (10/66) patients were diagnosed with biochemically silent PPGLs originating within parasympathetic paraganglia, invariably carotid body tumors. In contrast, patients in the second group, with childhood/early adult hypoxia onset, presented the opposite profile, with 64% (7/11) parasympathetic carotid body tumors and 36% (4/11) sympathetic tumors (**Fig. 4B, 4C, table S12)**. This result suggests for an early time window to the tumorigenic effect of hypoxia in the sympathetic system, possibly while the adrenal medulla and paraganglia are still uninnervated, developmental immature, and sensitive and autonomously responsive to low oxygen levels^17,18^

### Influence of oxygen levels on tumor development

We examined differences between patients with CCHD who developed PPGLs (N=149) and those who did not (N=1,599), delving into the analysis patients with reported SatO2 levels. The former group had significantly lower SatO_2_ than patients with CCHD without PPGLs (average SatO_2_: 81.2%, range: 68–91% vs. average SatO_2_: 83.2%, range 69–91%, respectively, p=0.01, **Fig. 4D, table S13**). Analyzing the subgroup of patients with CCHD who were not subjected to cardiac surgery, which more accurately represented the natural disease history, yielded similar results. Non-operated patients with CCHD who developed PPGLs had an average SatO_2_ of 79.3% (range: 68–90%), whereas non-operated patients with CCHD without PPGLs had an average SatO_2_ of 83.4% (range: 73–91%, p=0.0003, **Fig. 4E, table S13**). These data provide clinical support for the general but largely untested assumption that systemic hypoxia and different SatO_2_ levels could affect human tumorigenesis^93,94^.

### Influence of early hypoxia correction on tumor development

We observed a significant distinct surgical status within the CCHD cohort that developed PPGL tumors compared to those who did not. While 88% (1,204/1,368) of patients with CCHD without PPGL underwent early complete heart repair surgery, which indicated a return to normoxic conditions, only 17% (13/75) patients with CCHD who developed PPGLs underwent complete heart repair surgery (**Fig. 4F**, p < 0.001, **table S14**). Most patients (62/75, 83%) who developed PPGLs did not undergo surgery for heart disease, which implies that they maintained some degree of hypoxia throughout their lives or were offered only a palliative medical intervention that did not reverse hypoxia (**Fig. 4F, table S14**). This may suggest that early complete heart surgery that brought SatO_2_ to normal levels, could act as a protective factor against tumor development.

Altogether, this analysis revealed both tissue and time-dependency of hypoxia in tumorigenesis. Tissue dependency is related to the observation that patients with systemic hypoxia are enriched with tumors occurring exclusively in hypoxia-sensing and reacting cells, such as the carotid body, paraganglia, and adrenal medulla. Time dependency relates to hypoxia diagnosed since birth, favoring sympathetic PPGL development, whereas delayed hypoxia favors parasympathetic PPGL tumor (carotid body tumors) development. Our results also suggest that early surgical heart correction, which restores normoxia, may protect against PPGL development.

### Influence of systemic hypoxia and EPAS1-enriched PPGLs on long-term tumor features

To investigate systemic hypoxia effects on human tumor characteristics, we compared clinical features of 989 patients with PPGLs, with (N = 149, CCHD-PPGLs) or without (N =840, non-CCHD-PPGLs) congenital systemic hypoxia. Although the formal genetic status was not always determined to this extended cohort, based on the high *EPAS1* mutation frequency identified in our cohort with tumor samples available (**Fig. 1A**), most patients with CCHD-PPGL were assumed to have *EPAS1*^MUT^ tumors. Compared to patients with PPGL but without systemic hypoxia, those with CCHD-PPGL were younger at the time of PPGL diagnosis (Odds ratio, OR = 0.92; Confidence interval, CI: 0.91–0.94; p < 0.001), presented more frequently with paragangliomas than pheochromocytomas (OR = 9.98, CI: 6.19–16.11, p < 0.001), and were more likely to have multiple PPGL tumors (OR = 1.99, CI: 1.17–3.25, p = 0.008). Although PPGLs are usually non-metastatic tumors, patients with CCHD were at substantially increased risk of developing metastatic disease compared with those without CCHD who were diagnosed with PPGL (OR = 2.34, CI: 1.11–4.59, p = 0.018) (**Fig. 5A, 5B, table S15-S16**). These results indicate that systemic hypoxia increases the risk of developing PPGLs, and modulates tumor phenotypes, being associated with greater aggressiveness and metastatic risk.

**Fig. 5.**
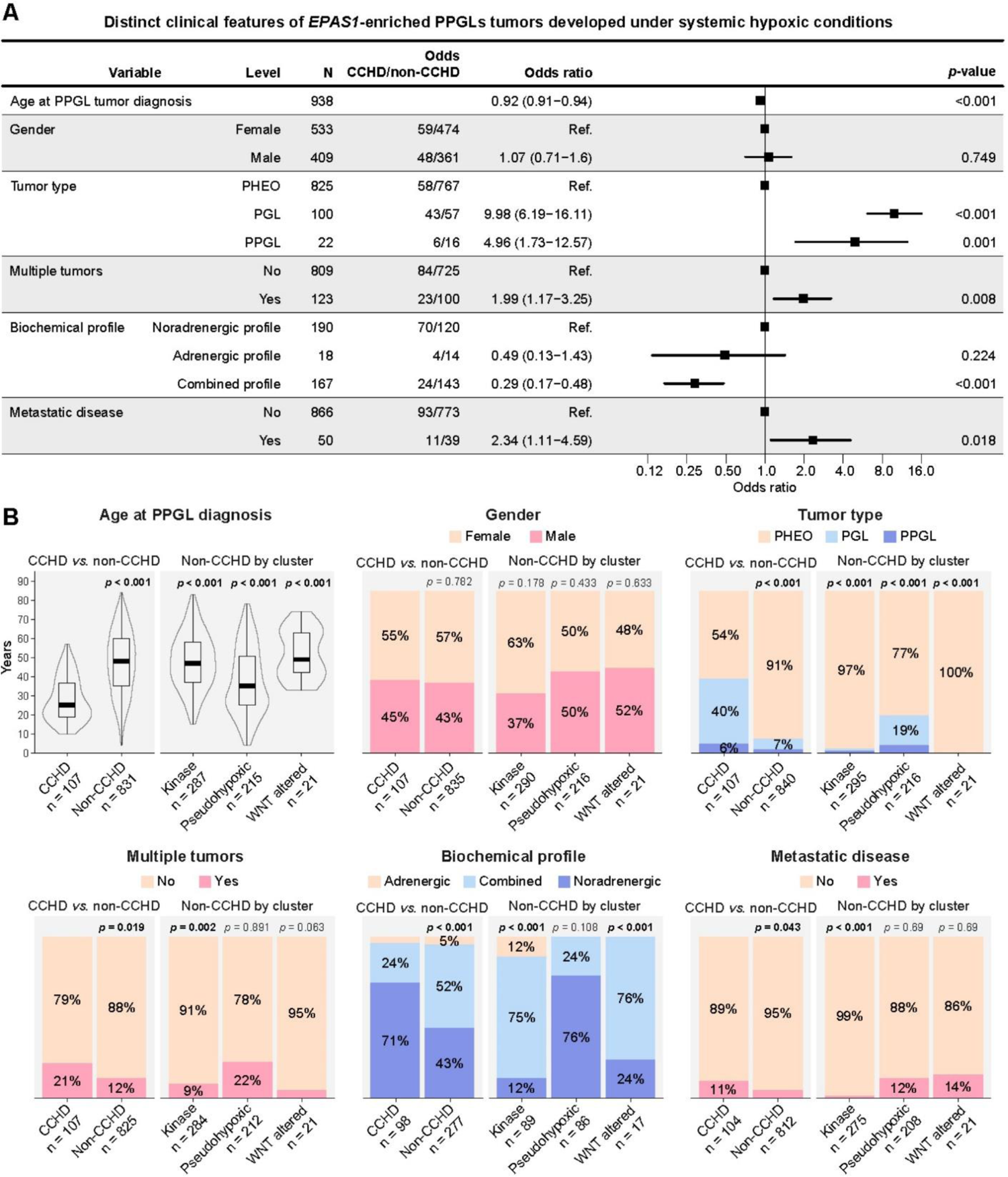
Hypoxic impact in PPGL tumor development: clinical features comparison of patients with CCHD-PPGL and without CCHD-PPGL (non-CCHD-PPGL). **A**) Univariate regression model (ODDs Ratio) of patients with PPGL and CCHD-PPGL. **B**) Representation of clinical variables in patients with PPGL and CCHD-PPGL. Cyanotic congenital heart disease (CCHD); Pheochromocytomas and paragangliomas (PPGL).

## Discussion

Our study presents *EPAS1*^HIF2α^ as a fundamental driver of evolutionary adaptation to systemic hypoxia, evident across extremely diverse conditions such as in populations living in high-altitude hypoxia^22–30^ and during human tumor development under early and prolonged systemic hypoxia (**Fig. 1D**). The findings of this study broaden our knowledge of the biological connectivity degree of life. This previously unreported broad convergence level suggests that other relevant molecular adaptation mechanisms occurring in natural populations may play a role in tumors facing similar stressors (**fig. S11**). Hence, future projects could investigate, roles of genes selected in populations adapted to high UV in skin melanomas or the impact of genes selected in populations adapted to dietary restrictions (amino acids, fat, and carbohydrates) on tumor metabolism reprogramming. Overall, harnessing adaptation mechanisms in natural populations could open new avenues for enhancing the detection of cancer vulnerabilities and therapies.

Our study also describes various significant correspondences occurring in high-altitude populations and tumors developed in systemic hypoxia, such as accelerated adaptability, gene functional plasticity, and embryological and tissue-dependence effects of hypoxia stress.

### Accelerated adaptability

Accelerated adaptability mechanisms, such as increased mutability and advantageous genetic allele exchange between different species (adaptive introgression), have been observed in cell lines cultivated in hypoxia^80–84^, and in animals and human populations adapted to high-altitudes^35–38^. These processes culminate in a genetic pool to cope with the effects of low oxygen levels. We found that human tumors developed under hypoxia also underwent accelerated adaptability, with *EPAS1* mutations as tumorigenesis drivers. Our data suggest that this highly specific and multistep process is associated with decreased DNA repair in sympathetic nervous system cells, whereas healthy tissues surrounding PPGLs from patients with CCHD, such as fat, stroma, and adrenal cortex tissues, retain detectable DNA repair proteins (**Fig. 2A**). After this, sympathetic cells acquire genomic instability and somatic mutations, indicated by the increased tumor mutation burden (**Fig. 2D–F**). In this context, a mutation occurring in the *EPAS1*^HIF2a^ oxygen-sensing domain undergoes extreme positive selection (**fig. S9**). Our analysis of the tumor evolutionary history via calculating CCF localized *EPAS1* as a tumor-initiating mutation (trunk mutation) and an oncogenic driver of PPGL development under hypoxia (**Fig. 1C**). For unknown reasons, this process is restricted to sympathetic tissues (adrenal medulla and paraganglia) composed of cells still immature at birth, capable of sensing hypoxia, and promoting surges in catecholamine secretion^17,18,95–98^

### Positive selection of EPAS1 genetic variants exclusively in high-altitude and systemic hypoxia-developed tumors

We found that the positive selection of *EPAS1* genetic variants was exclusive to hypoxic conditions. They have reportedly undergone one of the strongest selections observed in highlander Tibetans and Sherpas with a 90% frequency and fixation index (*F*_st_) reaching 0.81^22,23,25^. In parallel, tumors developed under systemic hypoxia harbored > 90% of *EPAS1* mutations (**Fig. 1A**, **Fig. 1D**). This strong evolutionary process, known as *selective sweep*, has also been described in highlander populations across different species living at high altitudes, such as horses^26^, dogs ^27^, wolves^28^, pigs^29^, and ducks^30^. In contrast, *EPAS1* variants are infrequent or nonexistent in populations living at low altitudes or in tumors of patients with normoxia (**Fig. 1D**, **fig. S6-S7, table S6-S7**).

### Functional plasticity

Although the positive selection of high-altitude adaptation and tumor genomes developed under systemic hypoxia converge on *EPAS1*^HIF2α^, selected variants have distinct characteristics and confer opposite molecular consequences. Although Tibetans and Sherpas are enriched in non-coding variants located in the promoter region with a loss-of-function effect^31–33^, we showed that tumors developed under hypoxia carried specific missense mutations in the oxygen degradation domain (amino acids 529–542) that conferred resistance to VHL-dependent protein degradation, leading to gain-of-function and oncogenic effects^40,48,99^ (**fig. S12**). Missense mutations outside the oxygen degradation domain, that is, the Andean H194R variant, have also been shown to be inactivated and positively selected at high altitudes^34^. These differing effects uncover functional plasticity of HIF2α-related adaptations and survival to hypoxia.

Hypoxia is regulated by HIFs, comprising inducible and constitutive subunits. In normoxia, hydroxylation of specific prolines in HIFα units (Pro531 in HIF2a and Pro564 in HIF1a) signals oxygen sufficiency and causes VHL-dependent ubiquitination and proteasome-mediated degradation^7,9,100^. This hydroxylation does not occur in hypoxia, hence pleiotropically activating HIFα, mediated via transcriptional upregulation of numerous target genes involved in several pathways. HIF2α controls EPO expression^20,21^, which is important because while lowlander sojourners visiting high altitudes experience rapid increases in EPO and hemoglobin concentrations to compensate for acute hypoxia, high hemoglobin levels are detrimental in the long term because they cause blood viscosity that paradoxically causes poor oxygenation and recurrent abortions, thus decreasing reproductive fitness. Hence, *EPAS1* gene variants positively selected in humans and other animals from the Tibetan region hamper HIF2α activation, which decreases hypoxia response, reduces EPO production, and lowers hemoglobin concentration, enabling their survival and reproduction in high altitudes. Therefore, Tibetan women carrying *EPAS1*^LOF^ alleles present lower hemoglobin levels and a greater capacity to carry pregnancies to completion, explaining the observed selection sweep^101,102^ (**fig. S12**). The link between Tibetan *EPAS1*^LOF^ alleles and higher reproductive fitness under chronic hypoxia has been confirmed in genetically edited mice^103^.

Contrary to HIF2α-hampering genetic variants selected in highlanders, *EPAS1* mutations detected in PPGL tumors strongly activate HIF2α via affecting the HIF2α Pro531 hydroxylation site or neighboring residues at the oxygen-dependent degradation domain, impeding its VHL-mediated proteasome degradation^40,48,99^. We have previously shown that these *EPAS1*^HIF2α^-activating mutants are stable in normoxia, triggering expression of genes associated with cell differentiation and stemness, which increase the tumor growth capacity in xenografts^40^. Furthermore, these *EPAS1*^GOF^ mutants become further stabilized and activated when oxygen levels reduced from 21% to 1%, suggesting that they probably yielded additional survival advantages under hypoxia^40,48^. The 20-fold increase in *EPAS1*^GOF^ mutation frequency seen in normoxic PPGLs (non-CCHD-related) to hypoxic PPGLs (patients with CCHD, 4.5% to 90%, **Fig. 1D**) strongly suggests that *EPAS1*^GOF^ confers increased fitness to sympathetic chromaffin cells, which may favor tumor development (**fig. S12**).

### Time and tissue-dependent tumor risk of systemic hypoxia

Hypoxia is a prevalent environmental stressor contributing to morbidity and mortality in humans and marine life^104–107^. Adult-onset hypoxia is estimated to affect approximately a billion people with obstructive sleep apnea and 400 million people with chronic obstructive pulmonary disease worldwide^108^; at birth, hypoxia affects more than 80 million people living above 2,500 m^109^, and 1–2 infants per 1,000 live births have CCHD^110^. Hypoxia at birth has been linked to a six-fold increased risk of PPGLs, whereas no other tumor type is enriched to the same degree^54,55^. These observations are consistent with independent experimental data from William Kaelin and Peter Ratcliffe, who indicated that PPGL tumor development was promoted by defective adrenal development and developmental apoptosis^111,112^. Our results clearly show a distinct effect of systemic hypoxia when it starts from birth compared with afterwards during childhood. Patients with CCHD and systemic hypoxia since birth were enriched in sympathetic catecholamine-secreting tumors, consistent with the early impact of systemic hypoxia on developmentally immature cells from the adrenal medulla and paraganglia of babies. During human fetal development, the environment is characterized by significant hypoxemia, with O_2_ pressures as low as 20 mmHg^113^, lower than that encountered by climbers at the Everest summit at 8,850m (25–28 mmHg)^114,115^. It further decreases immediately after birth following umbilical cord clamping^95^ (**Fig. 4B**). At birth, immature chromaffin cells in the adrenal medulla sense an abrupt decline in oxygen levels, triggering an extraordinary surge in catecholamine production, reaching up to 100-fold higher than those in a resting adult^96,97^. This surge shields infants from hypoxia-induced harm and mortality, prompting essential physiological responses, including heightened breathing, heart rates, and cardiac output. These responses work in tandem with blood flow redirection to critical organs, such as the heart and brain. Consequently, healthy newborns attain normoxia within 3–5 min after birth^95^. However, infants born with CCHD continue to experience hypoxemia because the defective heart chambers contain a mixture of oxygenated and deoxygenated blood. At birth, chromaffin cells of adrenal medulla, which produce catecholamines, are still immature and oxygen-sensitive, which explains the postnatal catecholamine surge^18,98^. Approximately one week after birth, these cells mature, become innervated by splanchnic nerves, and lose their initial oxygen sensitivity.

In contrast, patients with CCHD and systemic hypoxia occurring later in childhood present proportionally fewer sympathetic lesions, in line with the time-specific oncogenic effect of systemic hypoxia and sympathetic PPGL tumor development. These patients are proportionally enriched with carotid body tumors, possibly the result of disrupted adaptation (or maladaptation) to systemic hypoxia exposure at any time during their lifespan, rather than a true tumorigenic process. The carotid body, sympathetic paraganglia, and adrenal medulla are dependent on HIF2α for their embryological development and physiological function in adults^14^ (**fig. S13**).

Patients with hypoxia and CCHD exclusively have a six-fold increased risk of developing PPGL tumors; the cause remains unknown. PPGLs are neural crest tumors derived from hypoxia-sensing and responsive cells of the adrenal medulla and sympathetic and parasympathetic paraganglia^14^. Our data showed that in addition to tumor type exclusivity, CCHD-PPGL is also a nearly genetically exclusive disease with *EPAS1* gene convergence. This clinical and genetic exclusiveness is probably a reflection of the unique embryological role and expression pattern of *EPAS1*^HIF2α^ itself^14,116,117^. While HIF1α appears early in evolution and is expressed ubiquitously, HIF2α arises later in evolution because of gene duplication, and its embryonic and adult gene expression remains primarily restricted to few cell types, but prominently autonomic lineage cells, that is, cells driving the normal development of the carotid body (parasympathetic) and sympathetic tissues, the organs that later will undergo tumorigenesis (**fig. S14**).

### Oxygen consumption and supply balance as an advantageous mechanism for survival and proliferation

Our results, obtained via oxygen consumption rate measurements in cell lines with constitutively expressed *EPAS1* mutations, transcriptomic profiling of *EPAS1*-mutant PPGLs, and rat cell lines exposed to hypoxia, offer compelling evidence that *EPAS1* modulates the ETC via regulating the atypical *COX4i2* isoform expression (**Fig. 3A**). These findings suggest that reduced ETC activity, which was previously observed *in vitro* under hypoxic conditions^88^, also occurs in tumors with *EPAS1* mutations that thrive in hypoxic environments. These observations suggest a potential mechanism that challenges the conventional understanding of *EPAS* ^MUT^ PPGL survival under low-oxygen conditions. Instead of enhancing the ETC to maximize utilization of all available oxygen molecules in hypoxia to produce 34 ATP molecules via mitochondrial respiration, our data suggest that the competitive edge in survival and proliferation observed in *EPAS1*^MUT^ tumors arises from efficient matching of the ETC with available oxygen supply (**Fig. 3A, fig. S10)**. This balance in the ETC and oxygen utilization, achieved via slowing down the ETC under hypoxic conditions, likely reduces the toxicity associated with an imbalance between ETC activity and oxygen supply, as observed in hypoxic and superoxic conditions. Similar findings regarding the beneficial effects of balancing oxygen consumption and supply have been reported in Leigh mitochondrial disease, which is caused by mutations that impair the ETC. When knockout mice for *Ndufs4*, a common cause of Leigh syndrome, were exposed to moderate hypoxia (11% O_2_), their survival rates were found to increase significantly, whereas moderate hyperoxia (55% O_2_) led to early deaths^118^. Hypoxia treatment also provided substantial protection and extended the survival of another Leigh-like mouse model with a *SDHC* homozygous knockout^119^. Heterozygous *SDHC* mutations predispose humans to PPGL^120^.

### Translation to the clinic

As a potential translation to the clinic, the observed dependency on HIF2α seen in CCHD-PPGLs could be therapeutically explored via treatment with a HIF2α inhibitor currently under investigation in clinical trials. Amongst them, belzutifan (also called PT-2977 or MK-6482) has recently been approved by the Food and Drug Administration for treating other HIF2α-driven tumors such as renal cell carcinoma, central nervous system hemangioblastomas, or pancreatic neuroendocrine tumors associated with VHL disease^43^. The clinical and molecular features of Pacak-Zhuang syndrome, a combination of polycythemia and PPGL, are distinctly similar to those of CCHD-PPGL. Recently, a patient with Pacak-Zhuang syndrome who developed polycythemia and multiple paragangliomas due to somatic mosaicism for an activating mutation in *EPAS1* similar to those identified in CCHD-PPGL tumors, had a remarkable rapid and sustained tumor response to belzutifan; the patient also showed resolution of hypertension and polycythemia^121^. Our study shows that CCHD-PPGL may also benefit from belzutifan treatment based on their shared HIF2α aberrancy, a therapeutically exploitable genetic driver^44^. Nevertheless, belzutifan remains untested as a therapeutic option in a select group of patients with CCHD-PPGL.

### Leveraging adaptation mechanisms in natural populations and tumors

The discovery that a single gene (*EPAS1*^HIF2α^) can confer benefits to both natural populations and tumors, enabling them to adapt and thrive under hypoxic conditions, raises the possibility of similar correlations existing in other scenarios. The genes involved in genetic adaptation to specific natural stressors may help cells overcome similar stressors during tumorigenesis, metastasis, and acquired therapeutic resistance. These aspects can be further explored in future studies.

## Conclusions

The genetic architecture of high-altitude populations is shaped by a strong positive selection of the *EPAS1* gene, encoding HIF2α^22–30,35,36^. We generated and integrated multi-layered hypoxia data from large clinical datasets, multi-omics analyses of tumors, and *in vitro* experiments. We found evidence of the highly specific effects of systemic hypoxia on tumorigenesis in a tissue, time, and oxygen level-dependent manners. Tumors from patients born with cardiac abnormalities resulting in chronic systemic hypoxemia mimic the genetic adaptation mechanisms observed in high-altitude dwellers harboring highly prevalent *EPAS1^HIF2α^* genetic variants. Mechanistically, our data support an intricate tumorigenesis model in which systemic hypoxia deregulates the DNA repair machinery of hypoxia-sensitive and responsive cells, thereby increasing their genetic pool of copy number alterations and mutations. In this state of genomic instability, we propose that *EPAS1^HIF2α^* mutations promote oxygen consumption rate optimization via expressing the atypical isoform *COX4i2*. This study provides clinical, evolutionary, and mechanistic insights into the effects of chronic hypoxia on tumorigenesis, and the unprecedented parallelism of genetic adaptations to hypoxia in two widely different contexts: in high-altitude dwellers and tumors developed under systemic hypoxia.

## Materials and Methods

### Experimental Design

In this study, we aimed to characterize the level of genetic adaptative parallelism amongst two very distant hypoxic conditions, such as high-altitude populations of the Himalayas (i.e. Sherpas and Tibetans) and hypoxia-sensing cells of the body that are prone to develop PPGL tumors when exposed to prolonged hypoxia due to a congenital blood-mixing heart defect (CCHD). We also aimed to characterize in deep the molecular mechanism underlying hypoxia-driven tumorigenesis in humans. To be able to tackle these goals, we generated and analysed multimodal data from clinically-informative hypoxic and normoxic patients; high- and low-altitude populations; genetic and genomic profiles of human tumors; status of DNA repair system in the tumors; bulk transcriptomic profiles of human tumors and cell lines cultivated in normoxia and hypoxia; single-cell transcriptomic profiles of the developmental trajectory of the hypoxia-sensing cells of the body; and *in vitro* experiments to assess the downstream effect of *EPAS1*^HIF2α^ in cellular respiration rate and oxygen balance. All the multimodal data generated and/or used in the study are available through publicly available biological repositories and/or provided as supplementary material.

### Ethical Committee

This study was conducted in accordance with the ethical principles of the Declaration of Helsinki and with the approval for research granted by the Ethics Committee from the Vall d’Hebron University Hospital (Barcelona, Spain) (PR(AG)158/2018); La Paz University Hospital (Madrid, Spain) (PI-3230); Central University Hospital of Asturias (Oviedo, Spain) (CEImPA 2020.547); Assistance Publique Hôpitaux de Paris (Paris, France) (#00011928); The University of Texas Health Science Center at San Antonio (San Antonio, TX, USA) (IRB #HSC06-069H.); Mayo Clinic (Rochester, MN, USA) (14-008336); Brigham and Women’s Hospital, Harvard Medical School (Boston, USA) (2013P000564); and others. The research was conducted in accordance with local data protection laws and all patients were provided with written informed consent. All data provided are anonymized in line with applicable laws and regulations.

### Cohort of CCHD-only, PPGL-only, and CCHD-PPGL patients

To generate the data for the analysis of the potential impact of systemic hypoxia on risk to develop tumor, on tumorigenesis, and on the modulation of phenotypic characteristics of the tumors, we needed large and informative cohorts of patients with and without PPGL tumors and with and without systemic hypoxia. To obtain such cohorts, we established a multidisciplinary International Consortium comprised of clinicians (cardiologists, endocrinologists, oncologists, radiologists), pathologists with expertise in endocrine tumors, genomic researchers, and molecular researchers from 39 Institutes from 5 countries, many of them involved in the research of PPGL and hypoxia field for decades. Clinical data were generated and curated from 2,854 patients with informative clinical conditions for the study, such as 1,599 patients with CCHD and not PPGL; 840 patients with sympathetic PPGL and not CCHD; 266 patients with parasympathetic PPGL (all carotid body tumors) and not CCHD (n=24^1^, n=52^2^ and sequenced data kindly shared by Robledo’s lab (n=190)); and 149 patients with CCHD-PPGL, which is a substantial number due to the rarity of the combined CCHD-PPGL condition. Detailed information of the patients is provided in **table S6**.

### Cardiac and endocrine features of the cohorts

The inclusion criteria for the CCHD cohorts were patients with cyanotic congenital heart disease with cyanosis defined as resting arterial oxygen saturation below 92%. Patients were divided by cardiac anatomy in six groups: functionally univentricular heart (UVH); transposition of the great arteries (TGA); tetralogy of Fallot (TOF) including pulmonary atresia with ventricular septal defect; pulmonary atresia with intact ventricular septum (PAIVS); ventricular septal defect (VSD); and others (**table S1**). According to the 32^nd^ Bethesda Conference classification^3^, 66.7% of patients had a CCHD of great complexity. More than half of the cohort was either unoperated or only underwent a palliative procedure, resulting in majority of patients who had been cyanotic for decades. None of the CCHD patients with PPGL had family history of PPGL-related hereditary syndromes. The diagnosis of pheochromocytoma and paraganglioma was confirmed by histologic confirmation and/or symptoms, altered biochemical parameters and imaging findings, as indicated by the guidelines^4^. The biochemical profile of the tumors was determined by standard catecholamine measurements in the plasma (**table S1**).

### Search for risk factors associated with tumor formation and tumor phenotypic variation

We searched whether clinical factors linked to hypoxia were potentially associated with increased risk to develop PPGL tumors and/or tumor aggressiveness. Using U-Mann-Whitney test, we compared the tumor type (sympathetic/parasympathetic lineage) developed in patients with congenital hypoxia (n=66) or infant/childhood hypoxia (n=11). We also compared the oxygen saturation levels and status of normoxia correction by heart surgery in CCHD patients that did not develop and those that did develop PPGL tumors (N=229 vs N=89 patients and N=1,368 vs N=75, respectively).

To test whether clinical factors linked to hypoxia were potentially modulating phenotypic characteristics of PPGL tumors, we performed a set of univariate binominal Generalized Lineal Models (GLM) to compare the following attributes: 1) age at PPGL diagnosis, 2) gender (two levels: Male and Female), 3) tumor type (three levels: PHEO, PGL, and PPGL), 4) multiplicity (two levels: Yes and No), 5) biochemical profile (three levels: Noradrenergic, Adrenergic, and Combined) and 6) metastasis (two levels: Yes and No) in patients that had developed PPGL tumors in a context of systemic normoxia (non-CCHD) or systemic hypoxic (CCHD). Similar analyses were performed separating the PPGLs by molecular clusters (pseudohypoxia, kinase and WNT cluster). As all analyses generated 24 different GLMs, results were correct for multiple testing and p-values were adjusted by controlling the false discovery rate^5^. The above-mentioned analyses are graphically illustrated in Fig. 5A-5B and fig. S15-S16.

### Statistical analysis

Percentages and frequencies are presented for qualitative variables and Chi-squared or Fisher test were used for comparisons. Means (standard deviations) and medians (25^th^ and 75^th^ percentiles) were calculated for quantitative variables, and U Mann-Whitney or Kruskal Wallis tests were used for comparisons depending on the number of groups. All statistical analyses were performed with the R software, Version 3.5.2 (R Core Team). GraphPad Prism 6 software was used for graphical presentation of the data.

### Biological samples from CCHD-PPGL and parasympathetic PPGL tumors

Biological samples for genetics analysis were obtained through the CCHD-PPGL International Consortium. In total, 27 patients diagnosed with CCHD (22 newly recruited patients and 5 from Vaidya, et al.^6^) developed 28 sympathetic PPGLs, including 14 pheochromocytomas (adrenal medulla), 14 thoracic-abdominal paragangliomas (6 retroperitoneal, 5 peri-aortic retroperitoneal, 2 aorto/retrocaval retroperitoneal and 1 mediastinal thoracic), and 9 parasympathetic head and neck paragangliomas, all carotid bodies (CBs). Eighteen patients presented with a solitary tumor and seven with multiple and independent tumors (6 patients developed two tumors each, and one patient developed 5 independent tumors). P16 was the patient diagnosed with five PPGLs (1 mediastinal thoracic paraganglioma, 2 peri-aortic and 1 retrocaval abdominal paragangliomas and 1 pheochromocytoma, Fig. 1B). Five patients had synchronic tumors: two patients with pheochromocytoma + CB (P1, P12); one patient with bilateral pheochromocytoma (P#14); one with bilateral CB (P20); one with paraganglioma and pheochromocytoma (P26), and in two patients, the tumors were diagnosed during follow up: one patient with pheochromocytoma + paragangliomas + CB (P16) and other with bilateral pheochromocytoma (P21). P18 developed a hepatic metastasis. In addition, 60 parasympathetic carotid body^1^ (n=24) and head and neck (n=36) PPGLs developed in the context of systemic normoxia (non-CCHD) were obtained from The Central University Hospital of Asturias.

### Tumor and germline DNA extraction

Thirty-four of the 38 PPGL tumors (including a hepatic metastasis) from patients with CCHD were surgically excised and were available for genetic analysis (**table S1**). Tumor samples were obtained in formalin-fixed paraffin-embedded (FFPE) blocks from surgery and reviewed by a pathologist to confirm tumor type and the percentage of neoplastic cells was less than 20%, tumoral area was macrodissected prior DNA extraction using Maxwell 16 FFPE LEV DNA Purification Kit (Promega, Madison, WI, USA). DNA from the 60 non-CCHD carotid bodies were previously extracted and passed internal quality control at the VHIO’s laboratory before sequencing. The blood and saliva samples were collected using EDTA tubes and Oragene OGR-500 kit (DNA Genotek, Ottawa, Canada), respectively, and germline DNA was extracted using Gentra Puregene Blood Kit (Qiagen, Hilden, Germany). All DNA samples were quantified using a Qubit fluorometer device (Invitrogen, Carlsbad, CA, USA) before downstream genetic analyses.

### Genetic analysis

Genetic analyses on PPGL tumors from patients with CCHD and non-CCHD parasympathetic tumors were carried out using Sanger sequencing, next generation sequencing panel, and/or whole-exome sequencing (WES). All patients had tumor DNA sequenced and whenever available their paired gDNA were also sequenced (**table S1** for detailed information). For Sanger sequencing, the hotspot mutation region in exon 12 of the *EPAS1* gene was analyzed by PCR using previously described primers^7,8^: E12F-5’AACCCCCTTGCCTCTTTG and E12R-5’GGGGCAGATGGGGCTTAG, followed by capillary sequencing. Generated ABI files were used to assessment of *EPAS1* genetic status and variant allele frequency (VAF) using Mutation Surveyor software (SoftGenetics, PA, USA). For WES, 1000 ngs of leukocyte or tumor DNA were fragmented on a hydrodynamic shearing system (Covaris, Massachusetts, USA) to generate 180-280bp fragments. Remaining overhangs were converted into blunt ends via exonuclease/polymerase activities and enzymes were removed. After adenylation of 3’ ends of DNA fragments, adapter oligonucleotides were ligated. DNA fragments with ligated adapter molecules on both ends were selectively enriched in a PCR reaction. Whole exomic regions were captured using the SureSelect Human All V6 Exon kit (Agilent Technologies, CA, USA), purified using AMPure XP system (Beckman Coulter, Beverly, USA), and quantified using the Agilent high sensitivity DNA assay on the Agilent Bioanalyzer 2100 system. Sequencing was carried out on an Illumina HiSeq4000 platform with a median target coverage of ∼200X for tumor DNA and ∼100X for germline DNA.

Germline and tumor whole-exome sequencing data from 178 sympathetic PPGL tumors were retrieved from The Cancer Genome Atlas (TCGA)^9–11^. The newly generated WES from CCHD-PPGL patients and previously generated WES from the TCGA were processed as described below.

### WES bioinformatics analysis

Bioinformatics analyses were carried out following the GATK best practices^12^ and using pipelines described in our previous genomics study, code available in https://github.com/jfnavarro/scitron. The FASTQ files generated from the sequencers were turned into unaligned BAM files and adaptor sequences marked to generate uBAM files, which were aligned then to the hg38 reference using BWA. Read duplicates were marked and base quality scores recalibrated. Variant calling was carried out using Mutect2 with The Genome Aggregation Database (gnomAD)^13^ dataset as a population resource. The variants were filtered using information about normal contamination extracted from BAM pileups at the positions of Exac variants^14^, and from a strand orientation model. The following versions of the software were used: GATK (v4.1.9.0), Samtools (1.12), and BWA (0.7.17-r).

### Copy Number Variation, Purity, Ploidy, and Clonal structure of *EPAS1* mutations

For clonal structure analysis, the copy number variation of the segments containing the variant were calculated using ASCAT (version 3.1.1)^15^ as part of nf-core Sarek pipeline (version 3.3.2)^16–18^. ASCAT calculates allele-specific copy number changes in a tumor sample with respect to a matched normal sample. It also estimates the purity and ploidy of the sample. Cancer cell fraction (CCF) for *EPAS1* somatic mutations was calculated using the following formula^19^.

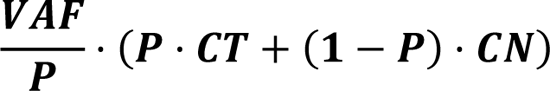

Where VAF is the variant allele frequency, P is the cellularity estimated from ASCAT, CT is the copy number of the segment covering the position in the tumor sample and CN is the same for the normal sample, which is assumed to be 2 except for autosomal chromosomes.

Clonal structure and phylogenetic reconstruction were performed using CONIPHER^20^ (version 2.1.0). Read support of autosomal SNVs and their associated copy numbers obtained from ASCAT were input to CONIPHER setting the parameter min_cluster_size = 2. Clonal distribution plots were constructed using the R package cloneMap^21^(version 1.0) using the CCF values calculated by CONIPHER for each clone.

### Analysis of Genomic Instability in PPGL tumors developed under systemic hypoxia or normoxia

Whole-exome sequencing data from 9 *EPAS1*-mutated sympathetic PPGL tumors was generated as described above. Whole-exome sequencing data from 170 sympathetic PPGL tumors from non-CCHD patients were obtained from Calsina B. *et al.*^22^. Briefly, the FASTQ sequencing files were aligned to the human reference genome GRCh37 using Burrows-Wheeler Aligner (BWA)^23^. Genome Analysis Toolkit GATK^24^, Haplotype Caller^25^ and MuTect^26^ were used to detect SNV and INDELs, and variant annotation was performed with Ensembl Variant Effect Predictor tool (VEP)^27^v90. SNV and INDELs were then manually curated as described in 1 to stablish the tumor mutational burden (TMB) of each tumor. Microsatellite instability (MSI) was assessed utilizing MANTIS^28^. Paired normal-tumor BAM files were aligned and processed, enabling the determination of microsatellite distribution discrepancies at the somatic level. Somatic copy number (CN) detection was performed with FACETS^29^. The total count of copy number (CN) segments, defined as SCNA burden, was computed for each sample. Details on the bioinformatic analysis of genomic instability can be obtained in Calsina B *et al.*^22^.

### Analysis of frequency of *EPAS1* somatic mutations in across human tumor types (non-PPGL)

We interrogated the *EPAS1* somatic mutation status in genomic data of 10,967 tumor samples from 32 different cancer types available through the PanCancer Atlas Studies of The Cancer Genome Atlas (TCGA) on the cBioPortal platform (http://www.cbioportal.org)^9–11^. Considering previous functional studies on *EPAS1* genetic variants^7,30^, only those occurring within the oxygen-dependent degradation domain of HIF2α (amino acid residues from 528 to 542) were considered to be activating. The *EPAS1* status of the nearly 11,000 human tumors analyzed is provided in **table S6.**

### Selection inference of *EPAS1* mutations across human tumors considering normoxia or hypoxia conditions

To characterize specificity of *EPAS1* mutations according to tumor type and hypoxia condition, we inferred positive selection of these mutations on the WES data generated for CCHD-PPGL tumors and over 10,000 tumors from 33 tumor types in the TCGA collection. Somatic mutation calls for primary tumors from TCGA were downloaded from the MC3 database^31^. We inferred selection on protein-coding genes with the Cancer Bayesian Selection Estimation (CBaSE) tool^32^ separately for three cohorts: (1) 184 PPGLs from TCGA (putative non-CCHD-PPGLs)^33^, (2) 10106 TCGA tumours from 32 non-PPGL tumor types, and (3) 14 tumours from CCHD-PPGL patients from this study. Since the CCHD-PPGL tumour samples were stored in FFPE, we manually filtered and curated the mutations as previously reported^22^, leaving a total of 341 short insertions and deletions, out of which 301 were coding single-nucleotide variants (i.e., missense, nonsense, or synonymous variants). *EPAS1* gene had 8 missense mutations in the PPGL tumors from TCGA and 7 in the CCHD-PPGL cohort. q-values of mutations and mapped dN/dS are provided in table S2.

### Analysis of frequency of *EPAS1* genetic variants in low and high-altitude natural populations

Frequency of *EPAS1* genetic variants in 13 low-altitude populations (N=148,162 individuals) and 10 high-altitude populations (N=627 individuals) were retrieved from published data and integrated in table S6. High altitude populations consisted of eight populations living across The Himalayas (Sherpas, Tibetan highlanders, Thakali, Bumthang, Chali, Brokkat, Kurtöp, and Layap) and two populations from the Andes Mountain range in South America (Colla Andean from Argentina and Quechua highlanders from Peru). These high-altitude populations live in altitudes varying from 3,000m to 4,450m (average 3,749m of altitude) on an atmospheric pressure as low as 60% of the sea level value. As a reference, the atmospheric pressure at the summit of Everest (8,849m) is 30% of the sea level value. Frequency of *EPAS1* genetic variants in populations of non-human mammals as horse, pig, wolf, dog, and duck living in high altitude and their low-altitude counterpart population were also obtained from previous publication and integrated in table S6.

Nearly all *EPAS1* genetic variants found enriched in high altitude occur in non-coding 5’ regions of the gene, mostly at the gene promoter and enhancers of the gene and are provided in table S6.

Frequency of these *EPAS1* non-coding genetic variants were investigated in low-altitude populations closely related from the high-altitude populations (i.e. the Han population, which is the closest related population to the highlander Tibetans), as well as in hundreds of thousands of individuals from various low-altitude populations available through The Genome Aggregation Database (gnomAD) version 4, which is composed of 76,215 genomes from populations worldwide^13^. The frequencies of these populations can be found in table S7. Due to the large number of populations analyzed, Fig. 1D depicts the frequencies of *EPAS1* genetic variants in some of the low and high-altitude populations whereas full data can be accessible in table S6.

### Searching for downstream targets of *EPAS1*^HIF2α^ involved in hypoxia adaptation

As the protein encoded by *EPAS1* gene, HIF2α, is a transcription factor known to control the expression of hundreds of different genes, we explored RNAseq datasets to search for downstream targets potentially involved in the observed positive selection of the *EPAS1*-mutated clones. Briefly, we used the DESeq2 package (v.1.28.1)^34^ to determined differentially expressed genes in *EPAS1*^mutated^ compared to *EPAS1*^wild-type^ tumors in two independent PPGL datasets, one from the TCGA^33^ and the other from patients with hypoxia-associated haemoglobin disorders^35^. We also searched for differentially expressed genes in PPGL-derived PC12 cell line that we cultivated in different periods of short (6h, 12h, 24h, and 48h) or prolonged (36 days, approximately 860h) normoxia condition (37°C, 5% CO2 and 19-21% O_2_) and hypoxia condition (37°C, 5% CO2 and 1% O_2_ balanced with N_2_). Detailed information about the bioinformatics pipelines used for RNAseq analysis and about the culture conditions of the PC12 cells are shown in Supplementary Material.

### Measurements of DNA repair protein levels in the PPGL tumors

Immunohistochemistry was performed on an automatic Ventana Benchmark Ultra System using 4 µm sections of tumor FFPE blocks. Samples were stained with Hematoxilin and Eosin, with the specific positive control to neuroendocrine tumors (Chromogranin A, clone LK2H10), and the four markers to microsatellite instability (MLH1, clone M1; MSH2, clone G219-1129; MSH6, clone SP93; PMS2, clone A16-4). After deparaffinization of the sections, the labelling was performed using ULTRACC! antigen retrieval buffer for 64 min at 100° C (anti-MLH1 and anti-PMS2) or for 40 min at 100°C (anti-MSH2 and anti-MSH6), incubation with primary antibody at 36°C (52min, 16min, 20 min, and 40 min respectively) and detection with Ventana Optiview DAB. Immunohistochemistry results were scored by two pathologists from different institutions for nuclear positivity for MSI protein in three categories: positive (staining in >10% nuclear tumor cells), focal positivity (staining in 1%-10% nuclear tumor cells) and negative (staining in <1% nuclear tumor cells). H score was obtained by multiplying the pro-portion of cells showing nucleus staining and the intensity of staining (0, no staining; 1, weak; 2, moderate; and 3, strong).

### Microsatellite instability, Somatic Copy Number and Tumor Mutation Burden analyses

Exome sequencing data was processed as described in Calsina B. *et al*^22^. Briefly, the CCHD-sympathetic PPGL (sympPPGL) cohort was aligned to the human reference genome GRCh37 using Burrows-Wheeler Aligner (BWA)^23^. Genome Analysis Toolkit GATK^24^, Haplotype Caller^25^ and MuTect^26^ were used to detect SNV and INDELs, and variant annotation was performed with Ensembl Variant Effect Predictor tool (VEP)^27^v90. SNV and INDELs were then manually curated as described in 1 to stablish the tumor mutational burden (TMB) of each tumor.Microsatellite instability (MSI) was assessed utilizing MANTIS^28^. Paired normal-tumor BAM files were aligned and processed, enabling the determination of microsatellite distribution discrepancies at the somatic level. Somatic copy number (CN) detection was performed with FACETS^29^. The total count of copy number (CN) segments, defined as SCNA burden, was computed for each sample. The non-CCHD PPGL cohort data was obtained from^22^.

### Mitochondrial respiration in cells expressing mutant *EPAS1*^HIF2α^

Authenticated HEK293T cells were cultured in DMEM (Gibco) 10% fetal bovine serum supplemented with and 1% of Penicillin-Streptomycin (Sigma-Aldrich, St. Louis, MO, USA). Stable cell lines were generated according to a previously described protocol^7^. Briefly, cells were transfected using Lipofectamine 3000 transfection reagent (ThermoFisher Scientific, USA) and HIF2α WT and HIF2α P405A/P531A plasmids (Addgene plasmid IDs #26055 and #19006, respectively). Transfected cells were selected with 0.5-1.0 ug/ml puromycin, and stable expression efficiency were confirmed by Western Blot against HIF2α in different time points. Puromycin-resistant cells expressing the wild-type and mutant version of the HIF2α were used for experiments for mitochondrial respiration measurements.

Mitochondrial oxygen consumption rates (OCR) were measured using the Agilent Seahorse XF Cell Mito Stress Test Kit and Seahorse XFe96 FluxPak (Agilent Technologies, 103010-100 and 103721-100) in the Seahorse XFe96 Analyzer (Agilent Technologies) following the manufacturer’s instructions. 3×10^4^ of HEK293T parental cells (condition 1), HEK293T stable cell line expressing HIF2α WT (condition 2), or HEK293T stable cell line expressing HIF2α P405/P531A (condition 3) were plated on poly-D-lysine coated plates on the day before of the assay including three replicate culture wells per run. Analysis was performed in the Seahorse XF DMEM Medium pH 7.4 (Agilent) supplemented with 10 mM glucose (Agilent), 2 mM glutamine (Gibco) and 1 mM sodium pyruvate (Gibco). Cells were washed twice with assay media and incubated for 1h in a 37°C non-CO2 incubator before starting the assay. Respiratory rates were measured in response to sequential injections of oligomycin (1.5 µM), FCCP (1.2 µM) and rotenone/antimycin A (0.5 µM) and were normalized to protein content per well using PierceTM BCA protein assay. The assays were performed twice with homogeneous results. The Seahorse Analytics software was used for analysis.

### *EPAS1* and *COX4i2* expression profiles in developmental and adult adrenal medulla

In previous studies, knockout of *EPAS1* deregulated catecholamine homeostasis and cardiac stimulus^36^ suggesting a direct effect of *EPAS1* on the embryological development of the sympathetic neural crest system and adrenal medulla, from which the PPGL tumors are derived from. To characterize the expression profiles of *EPAS1* and *COX4i2*, its downstream target involved in mitochondrial respiration, during maturation of neural crest derivatives, we analyzed single cell gene expression data of the entire neural crest lineage tree of mice embryos from the delamination to postnatal stages, including the domain of sympatho-adrenal differentiation. These data were obtained from a previous manuscript^37^ where the authors utilized deep sequencing method Smartseq2^38^ with the average of around 7000 genes per individual cell. Clustering and trajectory analysis was performed and gene expression plots of **Fig. 3D** were generated via Pagoda app (https://github.com/kharchenkolab/pagoda2).

### PPGL-derived PC12 cell lines cultivated in various normoxia and hypoxia conditions

PC12 cells derived from rat PPGL^39^ were cultured in normoxia condition (37°C, 5% CO_2_ and 21% O_2_) in DMEM medium supplemented with 15% v/v heat-inactivated fetal bovine serum, 100 U/ml penicillin, 200 µg/ml streptomycin and 2 mM L-glutamine (all from Gibco, Thermo Fisher Scientific, USA). Cells were then either continued cultivated in the same normoxia condition or were placed in a hypoxic incubator (HeraCell150, Heraeus, Hanau, Germany), that maintained a constant hypoxic environment (37°C, 5% CO2 and 1% O_2_ balanced with N_2_), for different short (6h, 12h, 24h, and 48h) or prolonged time (36 days, approximately 860h). Cell culture medium was changed whenever needed and all cells routinely tested and resulted negative for mycoplasma contamination.

Total RNA was isolated using the RNeasy mini kit (Qiagen, Germantown, MD), purity was checked using the NanoPhotometer® spectrophotometer (IMPLEN, CA, USA), and integrity and quantitation were assessed using the RNA Nano 6000 Assay Kit of the Bioanalyzer 2100 system (Agilent Technologies, CA, USA). 1μg RNA was used for sequencing libraries using NEBNext® UltraTM RNA Library Prep Kit for Illumina® (NEB, USA) following manufacturer’s recommendations and index codes were added to each sample. mRNA was purified using poly-T oligo-attached magnetic beads. After fragmentation with divalent cations in elevated temperature in NEBNext First Strand Synthesis Reaction Buffer (5X), first and second strand cDNAs were synthesized using random hexamer primer and M-MuLV Reverse Transcriptase (RNase H-) and DNA Polymerase I and RNase H, respectively. Remaining overhangs were converted into blunt ends via exonuclease/polymerase activities. After adenylation of 3’ ends of DNA fragments, NEBNext Adaptor with hairpin loop structure were ligated to prepare for hybridization. To select cDNA fragments of preferentially 150∼200 bp in length, the library fragments were purified with AMPure XP system (Beckman Coulter, Beverly, USA). Then 3 μl USER Enzyme (NEB, USA) was used with size-selected, adaptorligated cDNA at 37 °C for 15 min followed by 5 min at 95 °C before PCR. Then PCR was performed with Phusion High-Fidelity DNA polymerase, Universal PCR primers and Index Primer. PCR products were purified (AMPure XP system), library quality was assessed on the Agilent Bioanalyzer 2100 system and sequenced Illumina NovaSeq6000 sequencer.

### Analysis of differentially expressed genes in PC12 cells in hypoxia and PPGL tumors

For bioinformatics analysis, adapter removal and filtering of bad quality paired-end reads was performed using the *fastp* software (v.0.11.9)^40^ with the options: *-detect_adapter_for_pe --trim_poly_x --correction -r -M 10 -l 20*. The subsequent filtered reads were pseudo-aligned to the *rn6* rat genome and quantified at transcript-level using *Salmon* package (v.1.3.0)^41^. A prior salmon indexing step was created using a gentrome file composed by the concatenation of the Ensembl Rnor_6.0 transcriptome and genome assemblies. Transcript-level information was summarized to the gene-level for both exploratory and differential analyses with *tximport* package (v.1.16.1)^42^. The matrix of raw counts was rlog transformed for visualization purposes and normalized for DGE analyses according to *DESeq2* package (v.1.28.1)^34^. No-expressed genes (< 2 counts across all conditions) were discarded to reduce the multiple testing penalty. Likelihood ratio test (LRT) analysis was performed to identify differentially expressed genes between the three main conditions selected from exploratory studies (normoxia, hypoxia, and prolongued hypoxia, 36 days). Genes with an FDR<0.05 were considered as significant for downstream purposes. Gene co-expression analyses were performed using degPatterns function from DEGreport package (v.1.24.1) (http://lpantano.github.io/DEGreport/), on the rlog normalized expression matrix containing the combined significant genes detected across all comparisons in the aforementioned differential expression analyses. Over Representation Analysis of the biological functions associated with each of the modules was conducted with the goseq package (v.1.40.0)^43^ using Gene Ontology, KEGG and MSigDB Hallmark gene sets^44^. The gene length bias inherent to RNA-seq data was taken into account for these analyses. For the PPGL patient samples, log fold changes and q -values of genes were obtained from TCGA^9–11,33^. Differential expression analysis for the PPGL-derived cell line samples was performed using the R package DESeq2 (version1.40.0)^34^. Genes with adjusted p-value<0.01 were considered differentially expressed. The lfcShrink function from DESeq2 was used for visualization purposes. All gene expression data analyses were conducted using the statistical software R (v.4.0.2).

## Acknowledgments

We acknowledge Editage for editing the final versions of the text and figures provided by the authors. We acknowledge IdiPaz Biobank for the assistance in some tumor samples collection and the Oncology Data Science (ODysSey) Group of VHIO who carried out some statistical analyses. The authors would like to thank especially patients and their families for their participation and contribution to this research study, COST Action CA20122 Harmonisation for supportive networking. In memory of Dr. José Maria Oliver, who played a pivotal role in the development of the field of Adult Congenital Heart Diseases in Spain and his early assessment of CCHD-PPGL association. VHIO would like to acknowledge: the State Agency for Research (Agencia Estatal de Investigación) for the financial support as a Center of Excellence Severo Ochoa (CEX2020-001024-S/AEI/10.13039/501100011033), the Cellex Foundation for providing research facilities and equipment and the CERCA Programme from the Generalitat de Catalunya for their support on this research. The sponsors of this study listed below supported the costs related to the research experiments and salary of the investigators.

## Author Contributions

Conceptualization: RAT; Methodology: CA, JR-C, LC, BC, EG-G, DD, RF, ABM-C, JJA-L, BM, AMM-M, CA-E, BL, EAGG, SKF, EE, YD, RMRZ, JJP-K, CI, TD, CM, AOA, MEG-L-M, MJDC, IM-B, DML, MAAP, NB, AB, CB, PD, MFF, JLLP, JRS, FBF, SPAT, RC-B, RD, JC, APG-R, JF, DW, PN, WY, ARO, AV, IB, MR, AC, LD-S, MDC, IA, PLMD, RAT; Investigation: CA, LC, BC, EG-G, DD, RF, ABM-C, JJA-L, BM, AMM-M, SKF, EE, YD, TD, NB, AB, SPAT, DW, PN, AC, LD-S, MDC, IA, PLMD, RAT; Visualization: CA, BC, EG-G, DD, RF, DW, IA, RAT; Funding acquisition: RAT; Project administration: RAT; Supervision: PLMD, RAT; Writing – original draft: RAT; Writing – review & editing: CA, BC, EG-G, DW, ARO, LD-S, MDC, IA, PLMD, RAT.

## Data and Materials Availability

The clinical data of the patients included in the study (N=2,854), and the frequencies of *EPAS1* variants in natural populations (N=148,789) and in cancer cohorts (N=11,297) are available as supplementary material (**tables S1-S8**, **S12-S16**) for purposes of reproducing or extending the analysis. Newly generated whole-exome sequencing and RNAseq data (tables S9-S11) that support the findings of this study [will be] deposited in the European Genome Phenome Archive (EGA) with the accession codes [in progress].

## Supplementary Materials

Figs. S1 to S14

Tables S1 to S16

Material and Methods References

